# Predicting Sex-Specific Antiarrhythmic Strategies for Atrial Fibrillation through a Regression-Guided Computational Modeling Pipeline

**DOI:** 10.64898/2026.02.06.704432

**Authors:** Nathaniel T. Herrera, Haibo Ni, Charlotte E. R. Smith, Yixuan Wu, Dobromir Dobrev, Stefano Morotti, Eleonora Grandi

## Abstract

Atrial fibrillation (AF), the most common sustained cardiac arrhythmia, is a major contributor to stroke, heart failure, and mortality worldwide. Although AF affects both men and women at a similar rate, accumulating experimental and clinical evidence indicates that its underlying mechanisms, disease progression, and treatment responses differ by sex. However, current antiarrhythmic drug development and clinical management of AF remains largely sex neutral, likely contributing to limited efficacy and increased adverse effects. To address this gap, we developed a computational drug-screening pipeline based on experimentally constrained, sex-specific human atrial cardiomyocyte models to predict and evaluate sex-specific pharmacological strategies for AF. The pipeline integrates multivariable regression with mechanistic modeling to systematically test multi-target combinations of ion channel inhibitors and Ca^2+^ handling modulators and identify interventions that reduce arrhythmia vulnerability by restoring sex-specific electrophysiological and Ca^2+^ handling properties toward normal sinus rhythm (nSR). Application of this approach revealed a greater number of successful inhibitory drug combinations in males than in females. In males, optimal recovery to nSR primarily required inhibition of Na^+^ and K^+^ channels to prolong repolarization and refractoriness, increase Ca^2+^ transient amplitude (CaT_Amp_), and reduce susceptibility to action potential duration (APD) alternans. In females, modulation of Ca^2+^-related pathways was additionally required to suppress delayed afterdepolarizations (DADs). Forward single-cell simulations confirmed the predictions of the drug-analysis pipeline, demonstrating recovery of APD, CaT_Amp_, and arrhythmia vulnerability indices without introducing instabilities. Importantly, extension of these interventions to two-dimensional atrial tissue simulations demonstrated that sex-specific drug strategies reduce vulnerability to triggered activity, while suppression of reentry was most effective when combined with partial recovery of cell-cell coupling. Our results establish a multiscale computational pipeline for identifying sex-informed, multi-target antiarrhythmic therapies, amenable to experimental validation and translation to the clinic.

**Clinical Perspective:** *What is Known:*

- Atrial fibrillation arises from multiple interacting multiscale mechanisms, which limit the effectiveness of single-target therapies.
- Current single-target antiarrhythmic drugs for atrial fibrillation show more limited efficacy and higher adverse event rates in women than in men

*What the Study Adds:*

- This study demonstrates that effective pharmacologic strategies require different combinations of ion channel and calcium handling modulation in males versus females with persistent (chronic) atrial fibrillation.
- In males, coordinated Na^+^ and K^+^ channel inhibition most effectively improves electrical stability, whereas in females additional targeting of Ca^2+^ handling is required to suppress triggered activity.
- Sex-specific multi-target drug strategies including partial recovery of intercellular coupling suppress triggered activity and reentry in atrial tissue while preserving conduction.

## Introduction

Atrial fibrillation (AF), the most common sustained cardiac arrhythmia globally, affects over 33 million people and significantly contributes to adverse health outcomes, including ischemic stroke, heart failure, and premature death (Karatela & Calkins, 2025). Individuals with AF have a ∼50% higher risk of mortality and nearly double the risk for cardiovascular death (Odutayo et al., 2016). Importantly, arrhythmia susceptibility and clinical outcomes are profoundly influenced by sex (Ko et al., 2017; Odening et al., 2019). Males develop AF at younger ages, whereas females present with a greater burden of comorbidities, more severe symptoms, higher thromboembolic risk, and a similar lifetime prevalence due to longer life expectancy (Ball et al., 2013; Essebag et al., 2007; Ko et al., 2017; Odening et al., 2019). These marked sex-specific disparities highlight the urgent need for effective precision therapeutic strategies to reduce AF incidence and improve outcomes across the population. Antiarrhythmic drugs (AADs) remain the primary pharmacological option for managing AF, serving as the foundation for both rate and rhythm control. Despite their central role, AADs demonstrate only modest long-term efficacy and are limited by significant safety concerns, including proarrhythmic risk (Corley et al., 2004; Echt et al., 1991; Hassan et al., 2013; Saljic et al., 2023; Waldo et al., 1996). Notably, overall treatment efficacy in women is lower, with drug therapy more frequently discontinued due to adverse reactions, including pathological heart rate slowing when treated with amiodarone (Essebag et al., 2007; Muzzey et al., 2020; Rienstra et al., 2005). To address the limitations of conventional AADs, the HARMONY trial demonstrated that combining ranolazine with dronedarone significantly reduced AF burden relative to placebo and either agent alone, without additional safety concerns (Reiffel et al., 2015). Consistent with these findings, recent modeling studies have highlighted the promise of polytherapy approaches, in which coordinated block of multiple atrial-selective K^⁺^ currents could produce synergistic effects that enhance cardioversion and rhythm control in AF (Dasí et al., 2024; Ni et al., 2020).

Emerging mechanistic studies in human atrial tissue suggest that sex differences exist in the cellular and ionic substrates of AF arrhythmogenesis (Herraiz-Martínez et al., 2021; Odening et al., 2019; Smith et al., 2025). While these findings are derived from limited human cohorts or animal models and remain insufficient to establish definitive sex-specific mechanisms, computational modeling studies have begun to bridge this gap. We recently built experimentally constrained male and female human atrial cardiomyocyte models representing normal sinus rhythm (nSR) and chronic AF (cAF) to examine sex-dependent ionic and Ca^2+^ handling determinants of arrhythmic risk. We found that in cAF, males are more prone to action potential (AP) duration (APD) alternans, whereas females are more susceptible to Ca^2+^-driven triggers such as spontaneous Ca^2+^ release events and delayed afterdepolarizations (DADs; Herrera et al., 2025; Zhang et al., 2024).

Despite mounting evidence of these sex-based differences, AF therapies remain largely sex-agnostic, and no systematic approach exists to evaluate polytherapy strategies in a sex-specific manner, limiting translation of these mechanistic insights into precision therapeutic strategies. To address this gap, we developed a novel drug-screening in silico pipeline to identify sex-specific pharmacological strategies for AF. Using our populations of male and female human atrial cardiomyocyte models, we leveraged regression coefficients correlating changes in model parameters to key electrophysiological and Ca^2+^ handling biomarkers to systematically extract combinations of ion channel inhibitors (multi-channel block) capable of restoring electrophysiological and Ca^2+^ handling abnormalities in each sex. Predicted interventions were subsequently tested in forward single-cell simulations and extended to two-dimensional (2D) atrial tissue models to evaluate their impact on AP propagation, triggered activity, and reentry vulnerability. By integrating the synergistic effects of multi-target modulation with sex-dependent arrhythmogenic substrates across cellular and tissue scales, this study provides a framework for precision pharmacological strategies in AF. This multiscale approach supports the development of therapies that are both more effective and safer across sexes, and that can be directly evaluated in experimental and clinical electrophysiology settings.

## Methods

### Populations of Sex-Specific Atrial Myocyte Models

We used our recently developed and validated sex-specific human atrial cardiomyocyte models based on experimentally observed differences in ion channel and Ca^2+^ handling properties between males and females in nSR and cAF (Herrera et al., 2025, **Fig. 1**). For each sex, we generated a population of 1000 cAF model variants by perturbing the parameters listed in **Table 1** using log-normal scaling factors (σ = 0.1, Sobie, 2009). Model variants were paced to steady state at 1 Hz and evaluated using arrhythmia susceptibility protocols as previously described (Herrera et al., 2025). Briefly, five outputs were quantified for every model variant: i) APD at 90% repolarization (APD_90_) at 1 Hz, ii) Ca^2+^ transient (CaT) amplitude (CaT_Amp_) at 1 Hz, iii) maximum upstroke velocity (V_Max_) at 1 Hz, iv) alternans basic cycle length (BCL) threshold (Alt_BCL-T_), and v) DAD basic cycle length threshold (DAD_BCL-T_). Alternans thresholds were determined by progressively decreasing the BCL until beat-to-beat differences in APD estimated at *E*_m_ = -55 mV were ≥ 5 ms. DAD thresholds were evaluated following steady-state pacing at decreasing BCLs in the presence of 1 µM isoproterenol followed by a 30 s pause, with DADs defined as depolarizations ≥ 10 mV.

**Figure 1.**
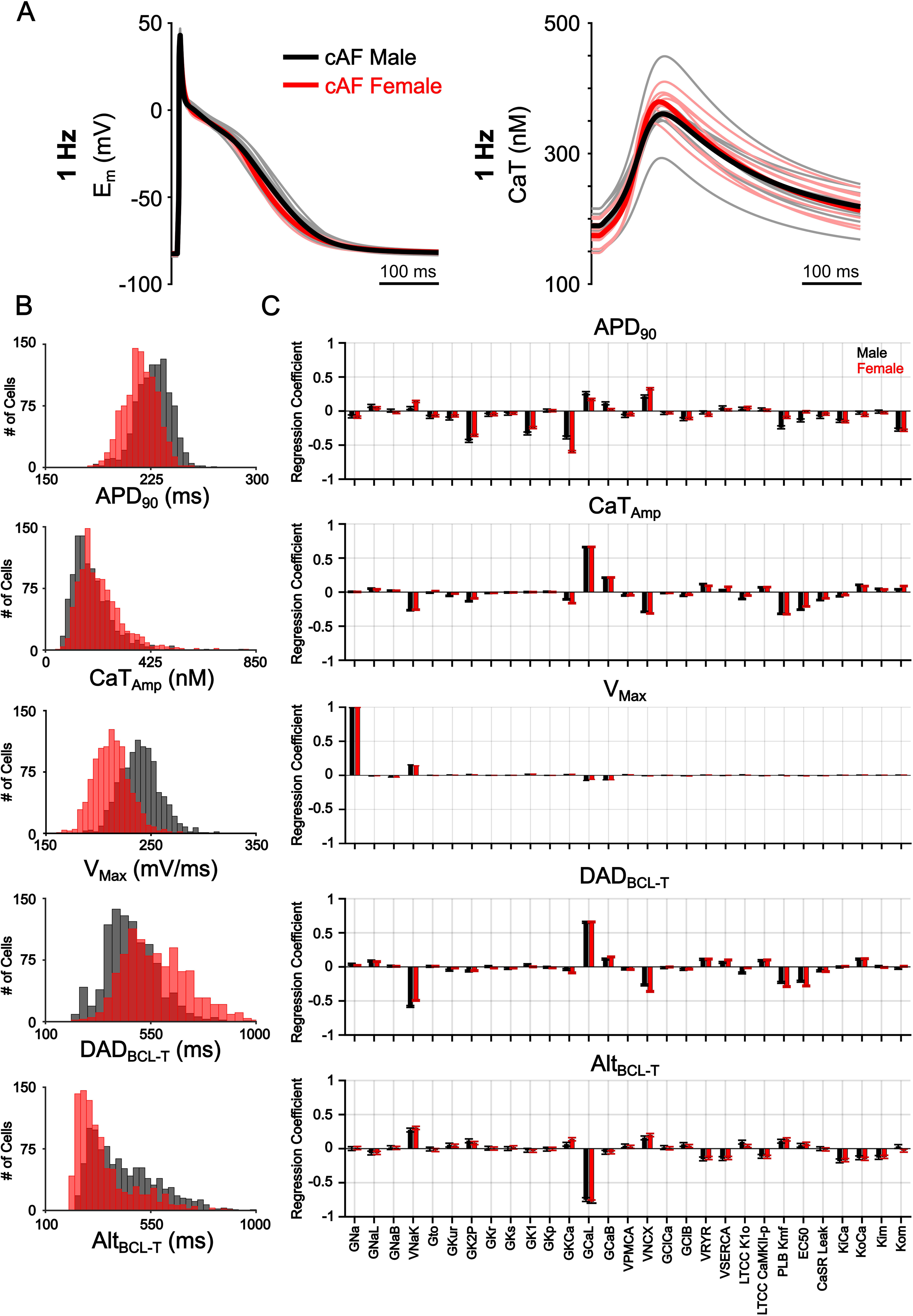
Population of sex-specific atrial cell models and regression coefficients in cAF. A) Human atrial action potentials (AP) and Ca^2+^ transients (CaT) at 1 Hz pacing from the validated male and female cAF baseline models, with 10 representative traces from each corresponding sex-specific model population (Herrera et al., 2025). B) Histogram distributions of each biomarker in the male and female cAF populations corresponding to the regression analyses shown in **panel C**. C) Multivariable regression coefficients for ionic and Ca^2+^ handling parameters (**Table 1**) quantifying the sensitivity of AP duration at 90% repolarization (APD_90_), Ca^2+^ transient amplitude (CaT_Amp_), maximum upstroke velocity (V_Max_), alternans basic cycle length threshold (Alt_BCL-T_), and delayed afterdepolarization basic cycle length threshold (DAD_BCL-T_), in male and female cAF model populations. Positive coefficients indicate that an increase in the parameter is associated with an increase in the biomarker, whereas negative coefficients indicate the opposite correlation. Error bars represent the 95% confidence intervals (CI), and horizontal ticks indicate the upper and lower CI boundaries.

**Table 1.**
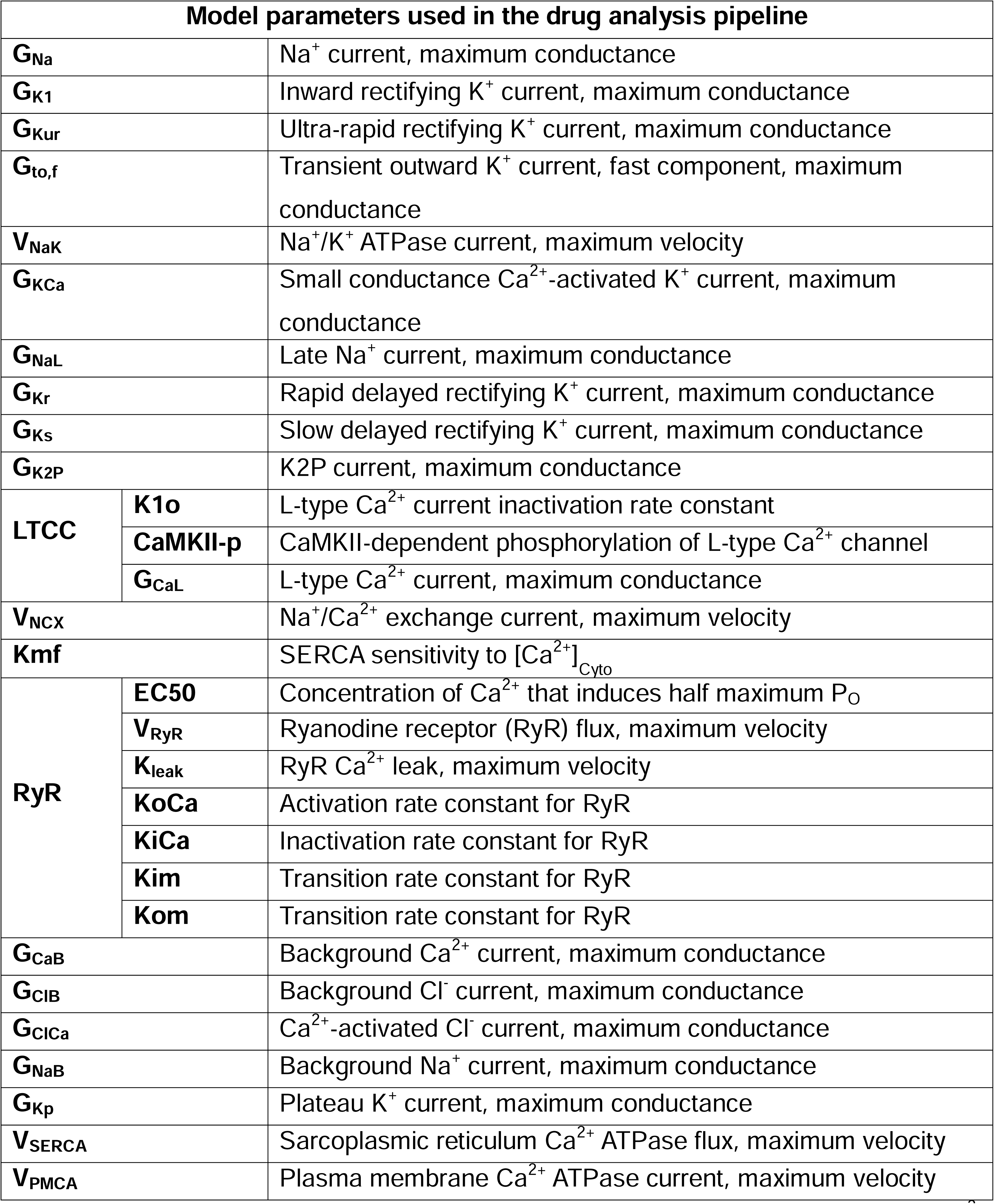
Model parameters used in drug analysis pipeline. Definition of ionic and Ca^2+^ handling parameters that are randomly varied to generate populations of models and are used in the linear regression-based pipeline to evaluate sex-specific drug effects and biomarker recovery.

### Multivariable Linear Regression

We performed multivariable linear regression analysis to correlate changes in model parameters with variations in each output (Sobie, 2009, **Fig. 1B and C**). For each sex, we constructed a regression matrix B that relates parameter perturbations to changes in model outputs such that, Δy ≈ BΔp, where Δp represents the log-transformed and z-scored parameter scaling factors relative to the sex-specific cAF baseline model, and Δy represents the log-transformed and z-scored corresponding changes in the five biomarkers. The sensitivity coefficients in B indicate the directional influence of each parameter on each biomarker, whereby a positive coefficient indicates that increasing the parameter increases the output, while a negative coefficient indicates the opposite. For example, the predicted change in APD_90_ can be expressed as:

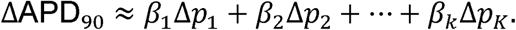

The full set of sex-specific regression coefficients for all five biomarkers are shown in **Fig. 1C**, which highlights distinct parameter-biomarker sensitivity profiles in male vs female in cAF.

### Biomarker Prediction and Parameter Inhibition Strategy

We next used the sex-specific regression matrices to systematically predict how concurrent modulation of multiple parameters (representing pharmacological inhibition of ion channels or Ca^2+^ handling targets) affect key electrophysiological biomarkers. Predicted biomarker changes (Δy) for a given parameter perturbation (Δp) were calculated using the derived sex-specific regression coefficient matrix B, where Δy ≈ B Δp, here Δp denotes the applied parameter perturbations and Δy corresponds to the predicted changes in model outputs (e.g., APD_90_). For single-target inhibition of parameter i, where all other parameters remain unchanged, the predicted change in APD_90_ for example is:

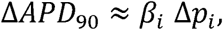

Each parameter was reduced using scaling factors from 0.90 (10% inhibition) to 0.50 (50% inhibition) and the analysis was expanded to multi-target interventions by systematically combining inhibitory effects across several parameters (**Fig. 2A**). For example, a 2-target (2-cross) combination involving parameters i and j, the predicted change in APD_90_ is:

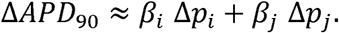

**Figure 2.**
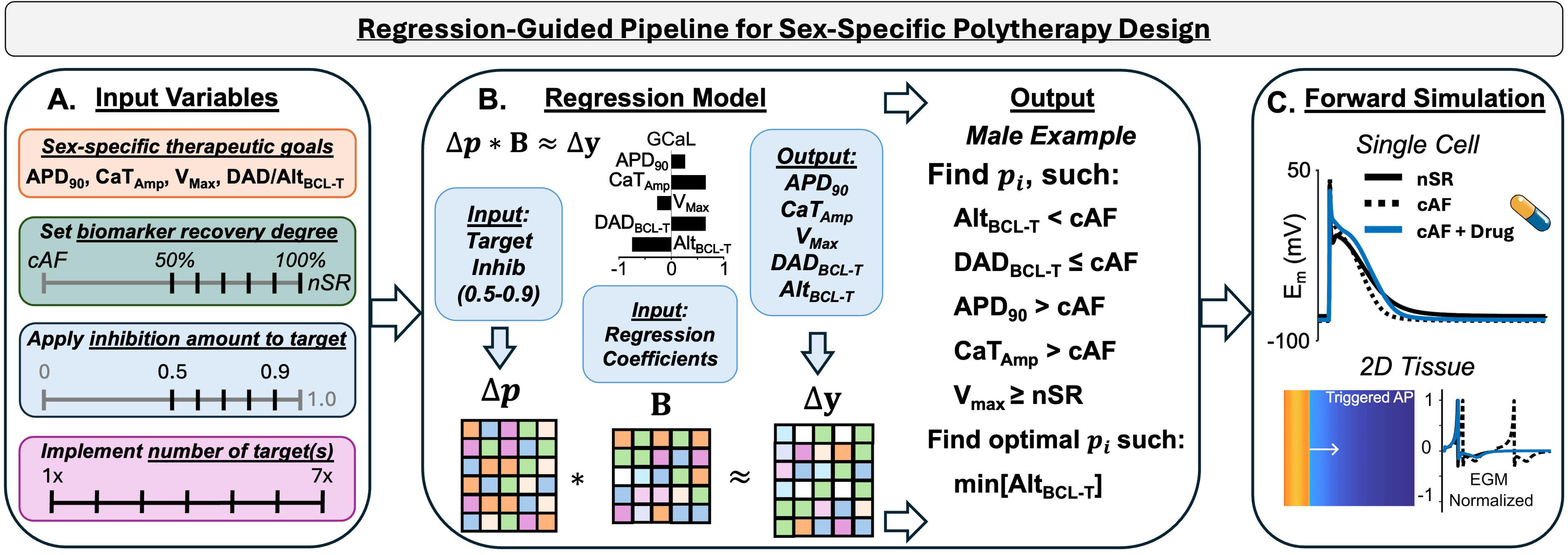
Regression-guided pipeline schematic. For illustration, the schematic is shown using the male case as an example. A) Input criteria for the regression-guided pipeline included restoration of APD_90_ and CaT_Amp_ toward nSR targets for both male and female, but the arrhythmia-specific metric differed (Alt_BCL-T_ in males; DAD_BCL-T_ in females). Recovery was evaluated across thresholds corresponding to 50-100% of the difference between the cAF and nSR values. Each parameter was independently scaled from 0.9x to 0.5x, and multi-parametric combinations of up to seven concurrent targets were tested to assess how combined inhibition influenced recovery across all biomarkers. B) Sex-specific multivariable linear regression was performed in male and female cAF model populations to relate parameter perturbations to changes in model outputs (APD_90_, CaT_Amp_, V_Max_, Alt_BCL-T_, and DAD_BCL-T_). Given the matrix of regression coefficients (B, Herrera et al., 2025) and specified inputs, sex (male/female), biomarker recovery (50-100%), target inhibition (0.5:0.1:0.9), number of targets (1-7), we computed the predicted outputs, and selected the optimal combination of parameter changes (p_i_) that minimizes arrhythmia risk by prioritizing reduction of Alt_BCL-T_ in males and DAD_BCL-T_ in females. These predicted perturbations were then applied in forward single-cell and 2D tissue simulations to quantify their effects on electrophysiological and Ca^2+^ handling biomarkers, as well as arrhythmia susceptibility in spatially coupled tissue. APD_90_, action potential duration at 90% repolarization; V_Max_, maximum upstroke velocity; CaT_Amp_, Ca^2+^ transient amplitude; Alt_BCL-T_, alternans basic cycle length threshold; DAD_BCL-T_, delayed afterdepolarization basic cycle length threshold.

The additive structure was similarly applied to higher-order combinations (3-cross, 4-cross, etc.).

### Go/No-Go Criteria

The goal of our screening pipeline is to identify combinations of parameter changes predicted to normalize abnormal cAF phenotypes toward the nSR ranges in each sex (**Table 2**). We established sex-specific recovery criteria across four biomarkers: APD_90_, CaT_Amp_, and the arrhythmia vulnerability metric relevant to each sex (Alt_BCL-T_ in males; DAD_BCL-T_ in females, **Fig. 2A**). To preserve physiological conduction properties, we additionally imposed a sex-specific constraint on maximal upstroke velocity (V_Max_), requiring that V_Max_ not fall below nSR values at 1 Hz pacing (≥ 205 mV/ms in females and ≥ 237 mV/ms in males), thereby preventing drug combinations that could impair conduction. To prevent unintended increases in arrhythmia vulnerability, we additionally required that DAD_BCL-T_ not increase relative to cAF in males, and Alt_BCL-T_ not increase relative to cAF in females. Recovery for each biomarker was evaluated across graded levels corresponding to 50% to 100% restoration of the difference between sex-specific cAF and nSR (**Fig. 2A**). An intervention was considered successful at a given recovery level only if all the required biomarkers met their respective target thresholds.

**Table 2.** Recovery Targets for Atrial Biomarkers. Electrophysiological and Ca^2+^ handling biomarkers used to quantify recovery toward sex-specific nSR values in the regression-guided drug analysis pipeline. Additional constraints were imposed on the maximum upstroke velocity (V_Max_) such that values could not fall below sex-specific nSR levels at 1 Hz (205 mV/ms in females; 237 mV/ms in males). APD_90_, action potential duration at 90% repolarization; CaT_Amp_, Ca^2+^ transient amplitude; Alt_BCL-T_, alternans basic cycle length threshold; DAD_BCL-T_, delayed afterdepolarization basic cycle length threshold.

| **Biomarker** | **Male Recovery** |  |  |  |  |  | **Female Recovery** |  |  |  |  |  |
| --- | --- | --- | --- | --- | --- | --- | --- | --- | --- | --- | --- | --- |
|  | **50%** | **60%** | **70%** | **80%** | **90%** | **100%** | **50%** | **60%** | **70%** | **80%** | **90%** | **100%** |
| **$APD_{90}$ (ms)** | 265 | 272 | 279 | 286 | 293 | 300 | 256 | 263 | 270 | 277 | 285 | 292 |
| **$CaT_{Amp}$ (nM)** | 215 | 224 | 233 | 242 | 251 | 260 | 239 | 247 | 254 | 261 | 268 | 275 |
| **$DAD_{BCL-T}$ (ms)** | 453 | 453 | 453 | 453 | 453 | 453 | 516 | 506 | 497 | 487 | 478 | 468 |
| **$Alt_{BCL-T}$ (ms)** | 327 | 312 | 297 | 282 | 267 | 252 | 305 | 305 | 305 | 305 | 305 | 305 |

### Utilizing the Regression-Guided Pipeline

For each pipeline execution, we specified: i) male or female sex, ii) the desired degree of biomarker recovery (50 to 100% restoration toward sex-specific nSR values), and iii) the number of parameter(s) to be inhibited within a given combination (1-7 targets). For the selected parameter(s), inhibition levels were systematically sampled from 0.90 to 0.50 in discrete steps of 0.1. This resulted in 29 single-target (1-cross), 406 two-target (2-cross), and 3,654 three-target (3-cross) combinations, with the number of possible inhibition profiles expanding to over 4.2 million when considering 7-cross combinations. For each combination of inhibition levels across the selected parameters, the regression coefficients (B) were used to calculate the predicted changes in APD_90_, CaT_Amp_, V_Max_, Alt_BCL-T_, and DAD_BCL-T_. Among combinations satisfying all recovery criteria, we selected the optimal set of perturbations for each drug combination using a sex-specific objective that minimized DAD_BCL-T_ in females and Alt_BCL-T_ in males (**Fig. 2B**). For each cross-size (target combination), we selected all the successful drugs that achieved the highest degree of recovery to apply in forward single-cell simulations (**Fig. 2C**).

### Polytherapy Testing via Single-Cell Forward Simulations

Sex-specific inhibitor combinations predicted by the analysis pipeline were applied to male and female cAF models, which were then paced to steady state at 1 Hz to quantify APD_90_, CaT_Amp_, and V_Max_ (**Fig. 4**). In addition, we evaluated the effects of each polytherapy strategy on arrhythmia susceptibility. To evaluate arrhythmia vulnerability, we applied the same arrhythmia susceptibility protocols described above for determining Alt_BCL-T_ and DAD_BCL-T_. From the set of drug combinations satisfying all recovery and safety constraints in single-cell forward simulations, we advanced a representative intervention per sex to tissue-level simulations. We chose the combinations achieving the highest recovery level toward sex-specific nSR values across the required biomarkers with the minimum number of targets.

### Polytherapy Testing via Two-Dimensional Tissue Simulations

A homogeneous 2D atrial tissue slab (225 x 200 cells) with a uniform spatial resolution of 0.25 mm was constructed using the sex-specific baseline single-cell atrial cardiomyocyte models, following previously described methods (Ni et al., 2023). To account for structural remodeling in cAF, intercellular coupling was uniformly reduced by 40% in both male and female tissue simulations to mimic the effects of increased fibrosis, resulting in physiologically relevant conduction velocity (CV) values at 1 Hz. Under nSR conditions, CV was 0.667 m/s in males and 0.636 m/s in females, while in cAF it was reduced to 0.467 m/s in males and 0.437 m/s in females (Silva Cunha et al., 2024). Electrograms (EGMs) were computed to quantify triggered activity in tissue simulations and were normalized to their maximal value.

### Simulation protocol for assessing triggered activity and reentry

To assess triggered activity, initial conditions for the 2D tissue were obtained from single-cell models paced to steady state at a BCL of 500 ms (2 Hz) in the presence of 0.1 µM isoproterenol. The tissue was paced using a 10-cell-wide (2.5 mm) stimulus band at the left boundary for three additional beats at 500 ms BCL, followed by a 3 s pause to allow spontaneous activity to develop. DADs were classified as triggered APs (tAPs) if the depolarization amplitude was ≥ 50 mV.

To assess reentry, initial conditions were obtained from single-cell models paced to steady state at a BCL of 1000 ms (1 Hz) and mapped to the 2D tissue. The tissue was paced on the left boundary similarly as before, for three S1 beats at 1000 ms, after which a cross-field S2 stimulus was applied to the lower band of 60 rows. The S1-S2 coupling interval was systematically varied (in 10 ms intervals) to identify the interval windows associated with propagation, wavebreak, reentry, and refractoriness.

### Numerical methods and code

Single-cell simulations, including the drug-analysis pipeline and forward simulations, were performed on a workstation with an Intel Core i9-10900X CPU at 3.70 GHz (10 cores) using MATLAB R2022a (MathWorks, Natick, MA). Tissue simulations were performed on a computing cluster with Intel Xeon CPU E5-2690 v4 at 2.60 GHz (28CPUs; 56 threads) with 132 GB RAM. Data analysis was conducted in MATLAB. The source code is available at https://github.com/drgrandilab.

## Results

### Sex-Specific Therapeutic Insights

Application of the regression-guided pipeline revealed distinct sex-dependent strategies required to restore human atrial electrophysiological and Ca^2+^ handling features toward nSR values in cAF. Interestingly, no single-channel inhibitors achieved the defined recovery criteria in either sex, thus highlighting the need for multi-target modulation. In males, a minimum of 4 simultaneous targets were required to achieve partial recovery (**Fig. 3A**), whereas in females at least 5 targets were needed (**Fig. 3B**). Across all target combinations and recovery levels, males had a greater number of successful combinations compared to females (**Fig. 3**). The identity of optimal targets also diverged between sexes. In males, the most effective combinations predominantly involved inhibition of Na^+^ and K^+^ currents (e.g., G_K2P_, G_KCa_, G_NaL_, **Fig. 3A**), whereas in females successful strategies required a broader inclusion of Ca^2+^-related targets (e.g., G_CaB_, L-Type Ca^2+^ channel, LTCC, inactivation rate K1o, and LTCC CaMKII-dependent phosphorylation, **Fig. 3B**) in addition to inhibition of Na⁺ and K⁺ currents. Additionally, successful recovery in both sexes was typically associated with moderate, rather than maximal (50%), inhibition of Ca^2+^ handling parameters, reflecting their increased sensitivity under the combined dynamic constraints imposed by DAD_BCL-T_ and CaT_Amp_ recovery (**Fig. 3B**).

**Figure 3.**
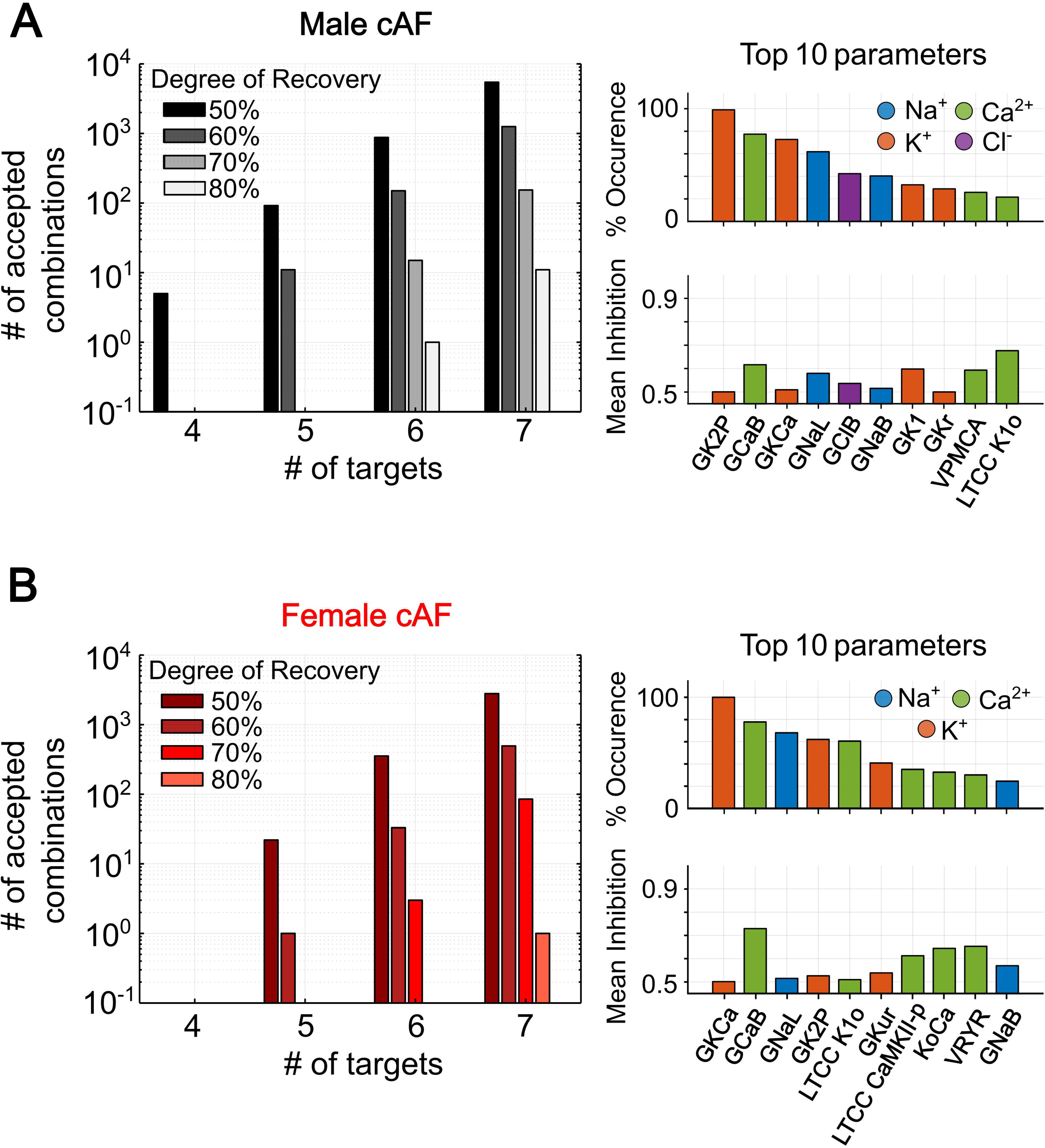
Regression-guided drug-analysis pipeline results for male and female cAF models. The left panel shows the distribution of accepted drug combinations as a function of the number of targets included in each intervention for male (A) and female (B). Accepted combinations were defined as those achieving biomarker recovery toward nSR values within prespecified thresholds. The right panel shows the ten most frequently selected parameters among all accepted combinations for male (A) and female (B), along with the average perturbation magnitude applied to each parameter. Colors denote the associated ionic processes

### Single-Cell Validation of Predicted Interventions in Forward Simulations

Forward simulations were performed to evaluate the multi-target interventions leading to the highest degree of recovery within each intervention size (i.e., number of concurrently inhibited targets), which led to 28 combinations in males and 5 in females (**Table 3**, **Fig. 4**). For each candidate therapy, we applied the optimal inhibition levels derived from the regression-guided pipeline for each parameter, and quantified APD_90_, CaT_Amp_, V_Max_, Alt_BCL-T_, and DAD_BCL-T_ (**Fig. 4**).

**Table 3.**
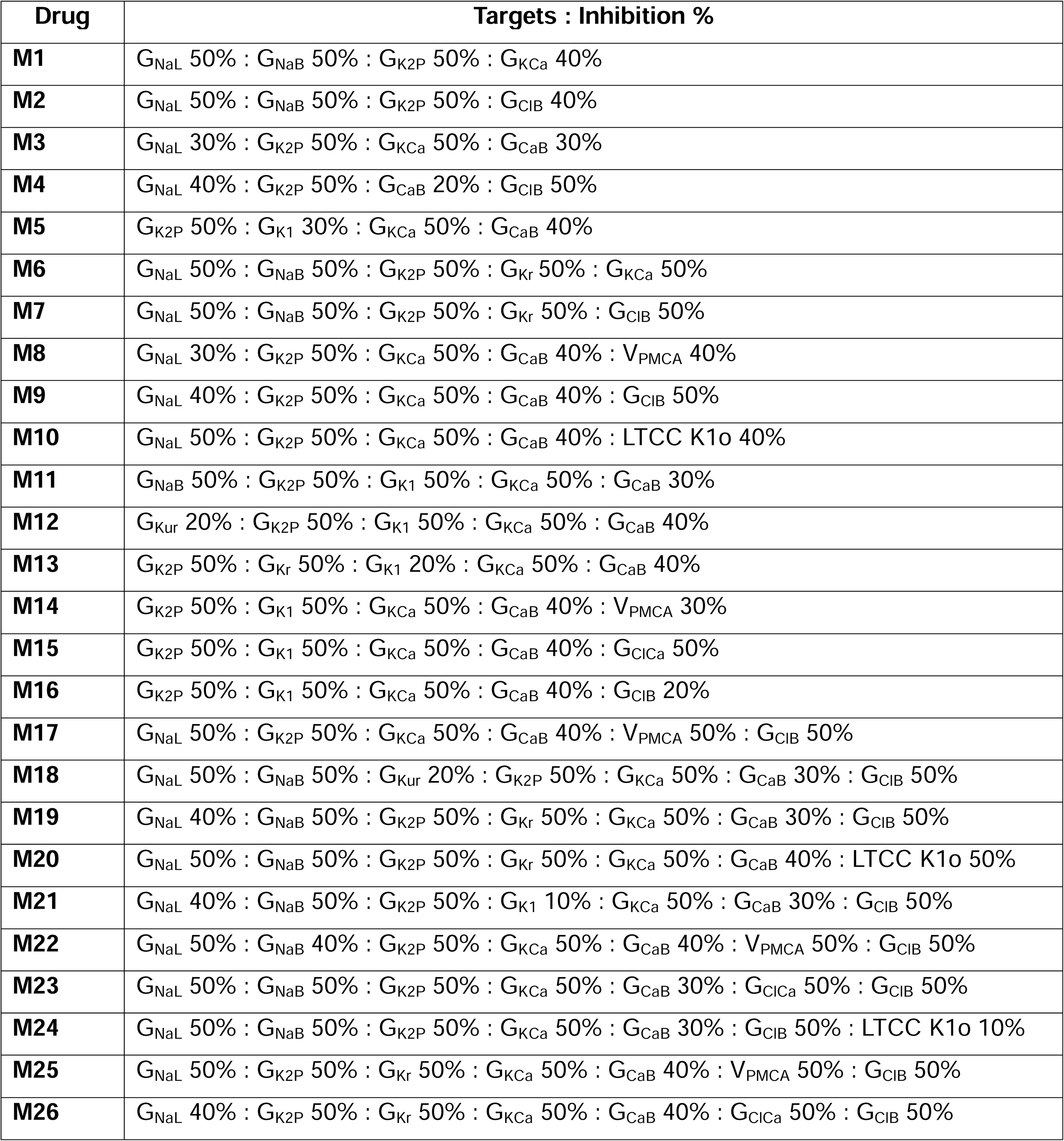
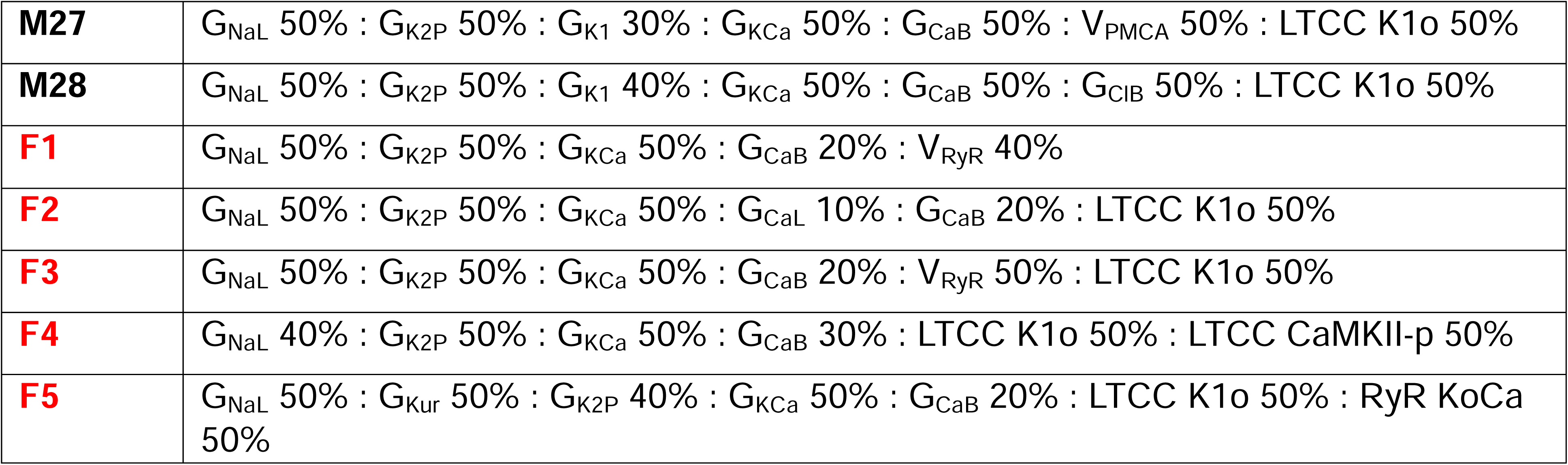
Multi-target parameter inhibition profiles used as simulated drug interventions in forward simulations. Each entry (M1-M28 for males; F1-F5 for females) denotes a regression-selected multi-target inhibition profile, where listed ionic and Ca^2+^ handling parameters are scaled to the indicated percentage of their sex-specific cAF baseline conductance (e.g., 40% = 0.60x baseline). These combinations correspond to the highest-recovery interventions within each target-number group identified by the regression-guided pipeline and were applied in forward single-cell simulations (**Fig. 4**).

| Drug | Targets : Inhibition % |
| --- | --- |
| **M1** | $G_{\text{NaL}}$ 50% : $G_{\text{NaB}}$ 50% : $G_{\text{K2P}}$ 50% : $G_{\text{KCa}}$ 40% |
| **M2** | $G_{\text{NaL}}$ 50% : $G_{\text{NaB}}$ 50% : $G_{\text{K2P}}$ 50% : $G_{\text{ClB}}$ 40% |
| **M3** | $G_{\text{NaL}}$ 30% : $G_{\text{K2P}}$ 50% : $G_{\text{KCa}}$ 50% : $G_{\text{CaB}}$ 30% |
| **M4** | $G_{\text{NaL}}$ 40% : $G_{\text{K2P}}$ 50% : $G_{\text{CaB}}$ 20% : $G_{\text{ClB}}$ 50% |
| **M5** | $G_{\text{K2P}}$ 50% : $G_{\text{K1}}$ 30% : $G_{\text{KCa}}$ 50% : $G_{\text{CaB}}$ 40% |
| **M6** | $G_{\text{NaL}}$ 50% : $G_{\text{NaB}}$ 50% : $G_{\text{K2P}}$ 50% : $G_{\text{Kr}}$ 50% : $G_{\text{KCa}}$ 50% |
| **M7** | $G_{\text{NaL}}$ 50% : $G_{\text{NaB}}$ 50% : $G_{\text{K2P}}$ 50% : $G_{\text{Kr}}$ 50% : $G_{\text{ClB}}$ 50% |
| **M8** | $G_{\text{NaL}}$ 30% : $G_{\text{K2P}}$ 50% : $G_{\text{KCa}}$ 50% : $G_{\text{CaB}}$ 40% : $V_{\text{PMCA}}$ 40% |
| **M9** | $G_{\text{NaL}}$ 40% : $G_{\text{K2P}}$ 50% : $G_{\text{KCa}}$ 50% : $G_{\text{CaB}}$ 40% : $G_{\text{ClB}}$ 50% |
| **M10** | $G_{\text{NaL}}$ 50% : $G_{\text{K2P}}$ 50% : $G_{\text{KCa}}$ 50% : $G_{\text{CaB}}$ 40% : LTCC K1o 40% |
| **M11** | $G_{\text{NaB}}$ 50% : $G_{\text{K2P}}$ 50% : $G_{\text{K1}}$ 50% : $G_{\text{KCa}}$ 50% : $G_{\text{CaB}}$ 30% |
| **M12** | $G_{\text{Kur}}$ 20% : $G_{\text{K2P}}$ 50% : $G_{\text{K1}}$ 50% : $G_{\text{KCa}}$ 50% : $G_{\text{CaB}}$ 40% |
| **M13** | $G_{\text{K2P}}$ 50% : $G_{\text{Kr}}$ 50% : $G_{\text{K1}}$ 20% : $G_{\text{KCa}}$ 50% : $G_{\text{CaB}}$ 40% |
| **M14** | $G_{\text{K2P}}$ 50% : $G_{\text{K1}}$ 50% : $G_{\text{KCa}}$ 50% : $G_{\text{CaB}}$ 40% : $V_{\text{PMCA}}$ 30% |
| **M15** | $G_{\text{K2P}}$ 50% : $G_{\text{K1}}$ 50% : $G_{\text{KCa}}$ 50% : $G_{\text{CaB}}$ 40% : $G_{\text{ClCa}}$ 50% |
| **M16** | $G_{\text{K2P}}$ 50% : $G_{\text{K1}}$ 50% : $G_{\text{KCa}}$ 50% : $G_{\text{CaB}}$ 40% : $G_{\text{ClB}}$ 20% |
| **M17** | $G_{\text{NaL}}$ 50% : $G_{\text{K2P}}$ 50% : $G_{\text{KCa}}$ 50% : $G_{\text{CaB}}$ 40% : $V_{\text{PMCA}}$ 50% : $G_{\text{ClB}}$ 50% |
| **M18** | $G_{\text{NaL}}$ 50% : $G_{\text{NaB}}$ 50% : $G_{\text{Kur}}$ 20% : $G_{\text{K2P}}$ 50% : $G_{\text{KCa}}$ 50% : $G_{\text{CaB}}$ 30% : $G_{\text{ClB}}$ 50% |
| **M19** | $G_{\text{NaL}}$ 40% : $G_{\text{NaB}}$ 50% : $G_{\text{K2P}}$ 50% : $G_{\text{Kr}}$ 50% : $G_{\text{KCa}}$ 50% : $G_{\text{CaB}}$ 30% : $G_{\text{ClB}}$ 50% |
| **M20** | $G_{\text{NaL}}$ 50% : $G_{\text{NaB}}$ 50% : $G_{\text{K2P}}$ 50% : $G_{\text{Kr}}$ 50% : $G_{\text{KCa}}$ 50% : $G_{\text{CaB}}$ 40% : LTCC K1o 50% |
| **M21** | $G_{\text{NaL}}$ 40% : $G_{\text{NaB}}$ 50% : $G_{\text{K2P}}$ 50% : $G_{\text{K1}}$ 10% : $G_{\text{KCa}}$ 50% : $G_{\text{CaB}}$ 30% : $G_{\text{ClB}}$ 50% |
| **M22** | $G_{\text{NaL}}$ 50% : $G_{\text{NaB}}$ 40% : $G_{\text{K2P}}$ 50% : $G_{\text{KCa}}$ 50% : $G_{\text{CaB}}$ 40% : $V_{\text{PMCA}}$ 50% : $G_{\text{ClB}}$ 50% |
| **M23** | $G_{\text{NaL}}$ 50% : $G_{\text{NaB}}$ 50% : $G_{\text{K2P}}$ 50% : $G_{\text{KCa}}$ 50% : $G_{\text{CaB}}$ 30% : $G_{\text{ClCa}}$ 50% : $G_{\text{ClB}}$ 50% |
| **M24** | $G_{\text{NaL}}$ 50% : $G_{\text{NaB}}$ 50% : $G_{\text{K2P}}$ 50% : $G_{\text{KCa}}$ 50% : $G_{\text{CaB}}$ 30% : $G_{\text{ClB}}$ 50% : LTCC K1o 10% |
| **M25** | $G_{\text{NaL}}$ 50% : $G_{\text{K2P}}$ 50% : $G_{\text{Kr}}$ 50% : $G_{\text{KCa}}$ 50% : $G_{\text{CaB}}$ 40% : $V_{\text{PMCA}}$ 50% : $G_{\text{ClB}}$ 50% |
| **M26** | $G_{\text{NaL}}$ 40% : $G_{\text{K2P}}$ 50% : $G_{\text{Kr}}$ 50% : $G_{\text{KCa}}$ 50% : $G_{\text{CaB}}$ 40% : $G_{\text{ClCa}}$ 50% : $G_{\text{ClB}}$ 50% |
| **M27** | $G_{NaL} 50\% : G_{K2P} 50\% : G_{K1} 30\% : G_{KCa} 50\% : G_{CaB} 50\% : V_{PMCA} 50\% : LTCC K1o 50\%$ |
| **M28** | $G_{NaL} 50\% : G_{K2P} 50\% : G_{K1} 40\% : G_{KCa} 50\% : G_{CaB} 50\% : G_{ClB} 50\% : LTCC K1o 50\%$ |
| **F1** | $G_{NaL} 50\% : G_{K2P} 50\% : G_{KCa} 50\% : G_{CaB} 20\% : V_{RyR} 40\%$ |
| **F2** | $G_{NaL} 50\% : G_{K2P} 50\% : G_{KCa} 50\% : G_{CaL} 10\% : G_{CaB} 20\% : LTCC K1o 50\%$ |
| **F3** | $G_{NaL} 50\% : G_{K2P} 50\% : G_{KCa} 50\% : G_{CaB} 20\% : V_{RyR} 50\% : LTCC K1o 50\%$ |
| **F4** | $G_{NaL} 40\% : G_{K2P} 50\% : G_{KCa} 50\% : G_{CaB} 30\% : LTCC K1o 50\% : LTCC CaMKII-p 50\%$ |
| **F5** | $G_{NaL} 50\% : G_{Kur} 50\% : G_{K2P} 40\% : G_{KCa} 50\% : G_{CaB} 20\% : LTCC K1o 50\% : RyR KoCa 50\%$ |

**Figure 4.**
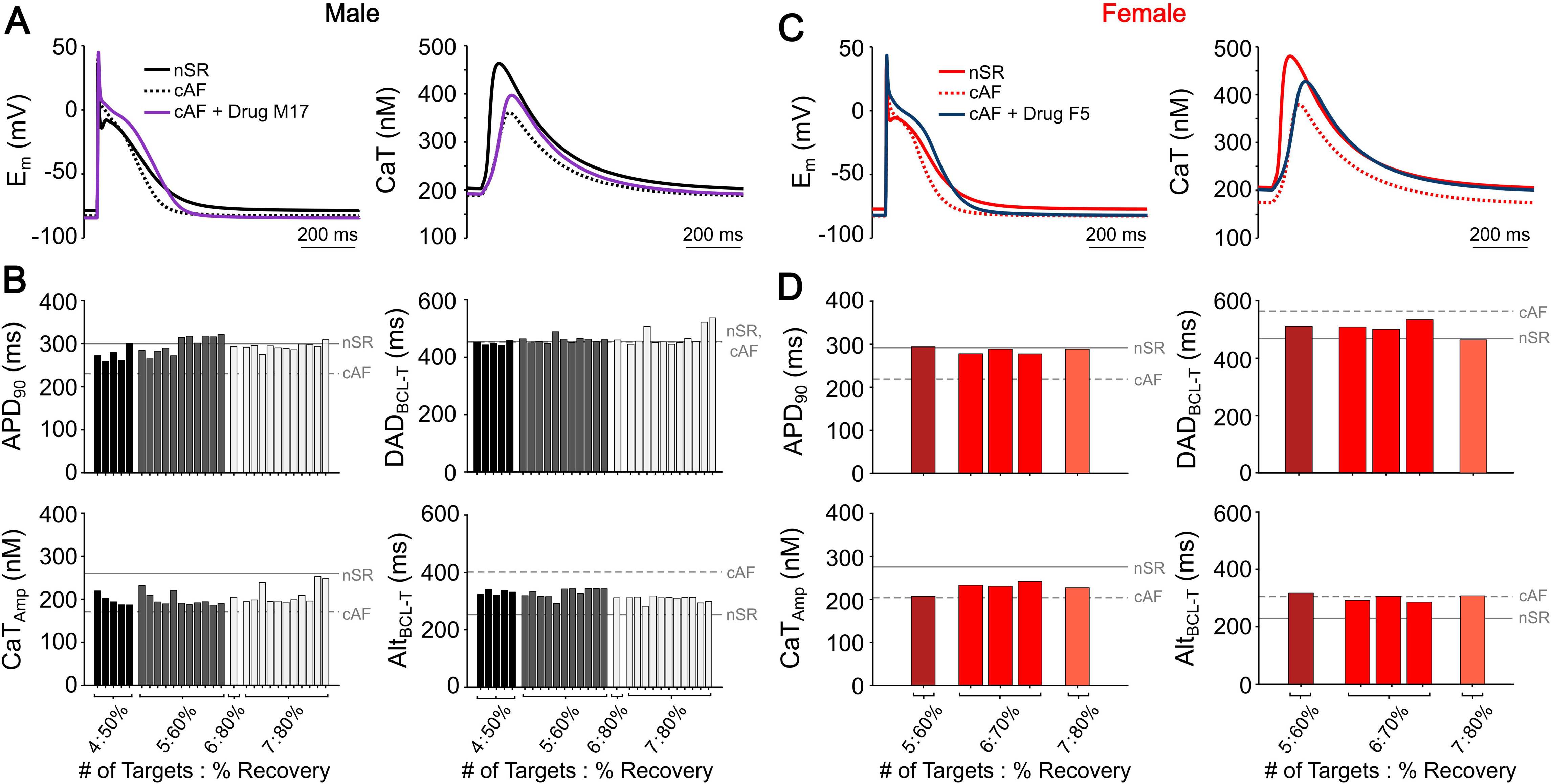
Forward single-cell validation of regression-guided drug interventions in male and female cAF models. A) Example of a successful intervention consisting of 6 targets achieving 80% recovery in male cAF. This drug corresponds to the 17^th^ bar (M17) in the plots shown in panel B below. Action potentials (APs) and calcium transient (CaT) are shown at 1 Hz pacing in nSR, cAF and cAF + M17. B) Forward-simulation results for the highest-recovery intervention at each target level in males. Effects are shown on APD_90_, CaT amplitude (CaT_Amp_), delayed afterdepolarization basic cycle length threshold (DAD_BCL-T_), and alternans basic cycle length threshold (Alt_BCL-T_). Solid grey lines indicate nSR single-cell reference values, and dotted grey lines indicate cAF baseline values. C) Example of a successful intervention consisting of 7 targets at 80% recovery in female cAF, corresponding to the 5^th^ bar (F5) in the plots shown in panel D below. AP and CaT traces are shown at 1 Hz pacing in nSR, cAF and cAF + F5. D) Forward-simulation results for female cAF interventions, showing effects on APD_90_, CaT_Amp_, DAD_BCL-T_, and Alt_BCL-T_, with grey solid and dotted lines indicating nSR single-cell reference and cAF baseline values respectively.

In males, all simulated interventions produced a longer APD_90_ and greater CaT_Amp_ at 1 Hz pacing (**Fig. 4A**). Increasing the number of targets yielded progressively greater biomarker recovery, with APD_90_ and CaT_Amp_ showing consistent improvement as additional targets were considered (**Fig. 4B**). Notably, V_Max_ remained above nSR values across all successful combinations. Importantly, Alt_BCL-T_ markedly improved with an increasing number of therapeutic targets, indicating that the predicted strategies could mitigate enhanced alternans susceptibility in cAF males (**Fig. 4B**). As intended by the design of the drug-analysis pipeline, DAD_BCL-T_ values remained stable across interventions (**Fig. 4B**), with only four exceptions (combinations M10, M20, M27 and M28; **Table 3**, **Fig. 4B**) exhibiting a slight increase in DAD_BCL-T_. In males, combination M17, which inhibits G_NaL_, G_K2P_, G_KCa_, G_CaB_, V_PMCA_, and G_ClB_ (**Table 3**), was selected for tissue-level evaluation because it achieved the greatest degree of recovery (80%) toward nSR with the least number of targets (6).

Female simulations similarly exhibited increased APD_90_ and CaT_Amp_ following drug application (**Fig. 4C**). Biomarker recovery improved with increasing target number, though to a lesser extent than in males (**Fig. 4D**). Across all drug combinations, APD_90_ exhibited robust recovery toward nSR values, CaT_Amp_ showed moderate to modest improvement, and V_Max_ was consistently maintained above nSR levels. In parallel, Alt_BCL-T_ did not increase, and DAD_BCL-T_ decreased as the number of targets in the combination increased (**Fig. 4D**). Notably, in females, combination F5, which inhibits G_NaL_, G_Kur_, G_K2P_, G_KCa_, G_CaB_, LTCC K1o, and RyR KoCa (**Table 3**), was chosen for tissue-level evaluation because it achieved 80% recovery toward nSR with the smallest number of targets (7).

### Tissue Validation of Predicted Interventions in Forward Simulations

While regression-guided screening and single-cell forward simulations identified sex-specific drug combinations that normalize electrophysiological and Ca^2+^ handling abnormalities and arrhythmogenic markers at the cellular level, we further sought to evaluate how cell-level sex-specific mechanisms and therapeutic interventions interact with tissue-level dynamics. We therefore evaluated predicted drug strategies in 2D atrial tissue to assess their effects on triggered activity and reentry vulnerability. To examine the interaction between pharmacological intervention and conduction, intercellular coupling was partially restored to a level consistent with the 80% recovery achieved for cellular electrophysiological and Ca^2+^ handling biomarkers under the selected sex-specific drug strategies.

Female nSR tissue exhibited stable behavior with no subthreshold DADs or tAPs, whereas cAF conditions displayed tAPs during the post-pacing pause. The optimized drug strategy (combination F5) was sufficient to suppress the triggered activity, both alone and when combined with improved conduction, whereas partial restoration of intercellular coupling alone did not abolish tAPs (**Fig. 5A**). In male 2D tissue paced at 2 Hz, neither nSR nor cAF conditions exhibited subthreshold DADs or tAPs during the pause period (**Fig. 5B**). Baseline stability was preserved with the optimized drug strategy (combination M17), with partial restoration of intercellular coupling alone, and when both interventions were simulated concomitantly (**Fig. 5B**).

**Figure 5.**
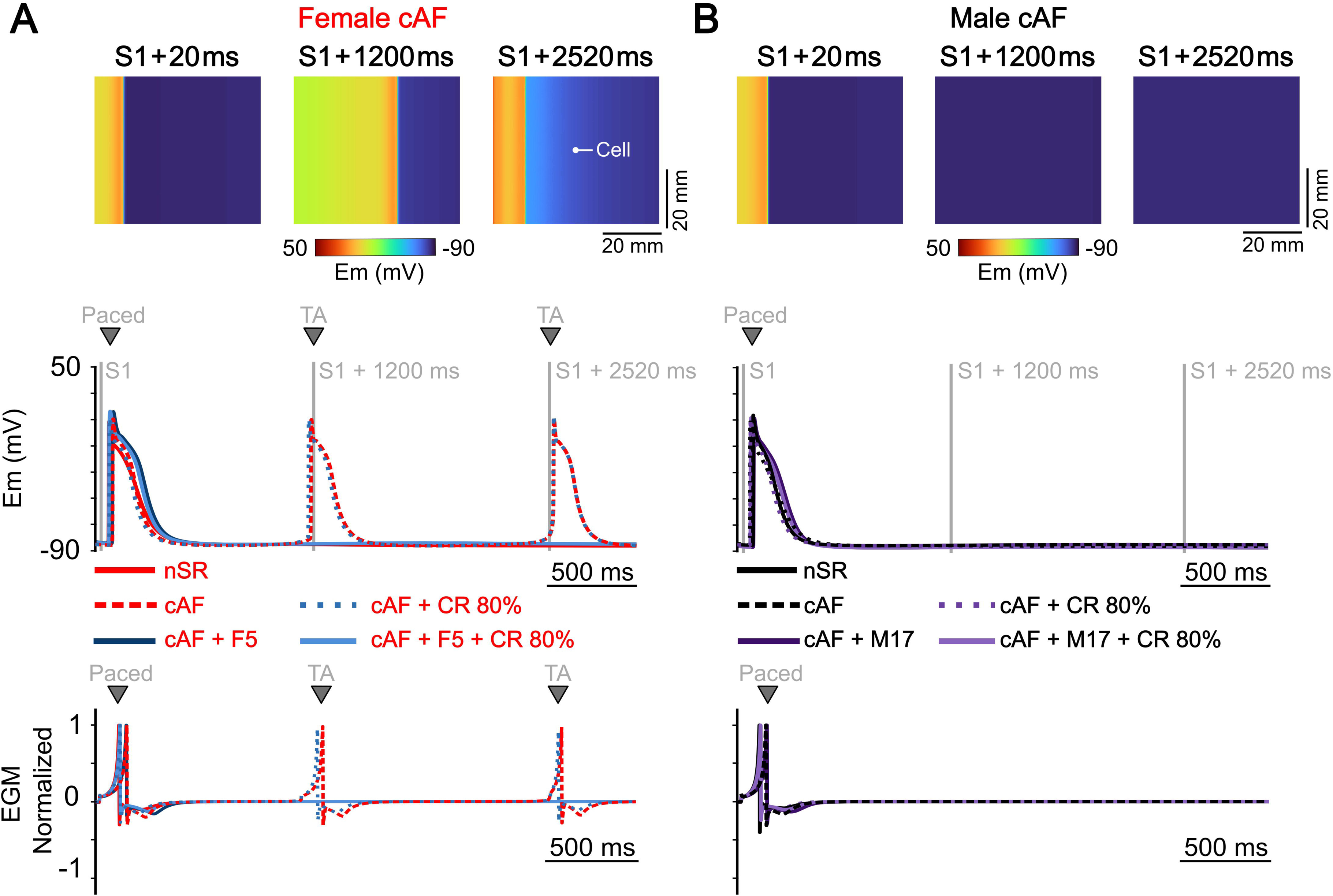
Sex-specific susceptibility to triggered activity and modulation by targeted interventions. The top row shows representative membrane voltage snapshots during the final S1 beat at 2 Hz in the presence of 0.1 µM isoproterenol, followed by the subsequent pause in female (A) and male (B). The middle row shows the corresponding single-cell action potentials (APs) extracted from the indicated tissue location under five conditions in female (A) and male (B): baseline nSR, cAF, sex-specific drug intervention, partial recovery of intercellular coupling (coupling recovery, CR), and the combined drug + CR. The bottom row shows the corresponding normalized electrograms (EGMs), in female (A) and male (B).

In male tissue, no reentry was observed in nSR, whereas sustained reentry over a vulnerable S2 window spanning 240-280 ms was exhibited in cAF. The optimized drug strategy (combination M17) narrowed the vulnerable window but shifted it to longer S2 intervals (300-310 ms). Partial restoration of intercellular coupling to 80% of nSR values similarly reduced the reentry window and shifted it to 240-260 ms. Most notably, combining M17 with coupling recovery abolished reentry across all tested S2 intervals (**Fig. 6C**). Importantly, CV at 1 Hz did not change beyond the AF-induced reduction, and increased only with partial recovery of intercellular coupling (nSR: 0.667 m/s; cAF: 0.467 m/s; M17: 0.444 m/s; coupling recovery: 0.596 m/s; M17 + coupling recovery: 0.583 m/s). Reentry did not occur in female tissue under nSR conditions but emerged in cAF over a vulnerable S2 window spanning 230-280 ms. The optimized female drug strategy (combination F5) substantially reduced reentry susceptibility but shifted the vulnerable window to longer S2 intervals (300-320 ms). Partial restoration of intercellular coupling to 80% of nSR values alone also narrowed and shifted the reentry window to 230-250 ms. Most importantly, the combined intervention of F5 with recovered coupling completely abolished reentry across all tested S2 intervals (**Fig. 6C**). As in males, CV at 1 Hz was not further reduced by pharmacological intervention in females, consistent with enforcement of the V_Max_ constraint (nSR: 0.636 m/s; cAF: 0.438 m/s; F5: 0.438 m/s; coupling recovery: 0.560 m/s; F5 + coupling recovery: 0.560 m/s).

**Figure 6.**
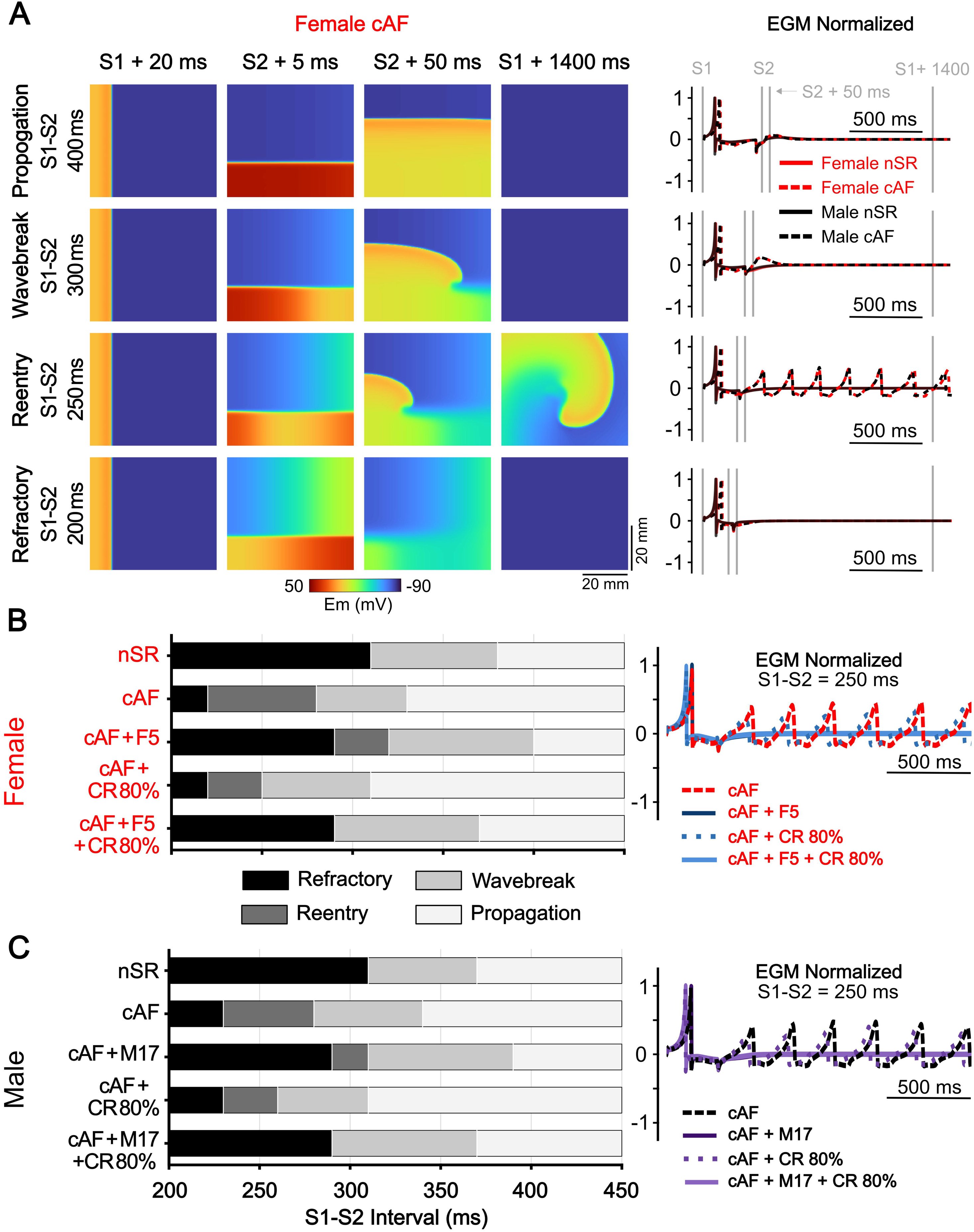
Sex-specific inducibility of reentry and modulation by targeted interventions. (A) Representative membrane voltage snapshots from female cAF atrial tissue illustrating the range of tissue responses to cross-field S2 stimulation. Four S1-S2 intervals are shown, selected to capture the four phenotypes observed in the inducibility protocol: propagation, wavebreak, sustained reentry, and refractoriness. For each S1-S2 interval, snapshots include the final S1 beat at BCL = 1000 ms, followed by three time points after S2 delivery (S2 + 5 ms, S2 + 50 ms, and S2 + 1400 ms) to visualize initiation and maintenance (or failure) of reentrant activity. The corresponding EGMs are shown to the right for baseline male and female tissue under nSR and cAF conditions. (B) Summary of reentry outcomes across S2 BCLs in female atrial tissue, classifying responses as propagation, wavebreak, reentry, or refractoriness. The panel to the right are EGMs at an S1-S2 interval of 250 ms, for four conditions: cAF, coupling recovery (CR), sex-specific drug intervention (F5), and CR + F5. (C) Summary of reentry outcomes across S1-S2 intervals in male atrial tissue, shown in the same format as in (B), with corresponding EGMs shown to the right.

## Discussion

Understanding how sex-specific remodeling shapes atrial electrophysiology and Ca^2+^ handling is critical for developing targeted antiarrhythmic therapies, especially because AF mechanisms are increasingly recognized to differ between males and females (Herraiz-Martínez et al., 2021; Herrera et al., 2025; Zhang et al., 2024). In this work, we used a regression-guided modeling pipeline to identify and validate sex-specific combinations of ionic and Ca^2+^ handling modulators that restore atrial cardiomyocyte function toward nSR. The results revealed that the therapeutic strategies required to achieve recovery differed between males and females, likely reflecting fundamental differences in the mechanisms driving arrhythmia vulnerability. Coordinated, multi-target interventions achieved more effective recovery than single-channel inhibition, demonstrating that the pathways contributing to electrical instability are both interconnected and sex-specific. Collectively, these findings highlight the need for therapeutic approaches that account for mechanistic differences to achieve safe and effective therapeutic strategies for AF.

### Use of regression for therapeutic intervention screening

Computational modeling has become a central tool for advancing our mechanistic understanding of AF, particularly through population-of-models and regression-based analyses that explicitly account for inter-subject variability (Ni et al., 2018). Using populations of human atrial models, prior studies have evaluated the electrophysiological consequences of AF-associated remodeling, demonstrating that altered ionic conductance’s and channel kinetics can drive APD shortening and repolarization changes (Sánchez et al., 2014; Zhu et al., 2021), that dysregulated Ca^2+^ handling promotes triggered activity through enhanced spontaneous Ca^2+^ release and DADs (Morotti & Grandi, 2017; Ni et al., 2023; Zhu et al., 2021), and at the tissue level, inter-subject variability in ionic current availability can influence reentry dynamics, conduction properties, and arrhythmia maintenance (Liberos et al., 2016). Building on these mechanistic insights, population-based computational frameworks have also been applied to evaluate anti-AF therapeutic interventions (Liberos et al., 2016; Sánchez et al., 2017), including multi-target modulation strategies, which have shown greater efficacy than single-target approaches in both experimental models and clinical studies (Dasí et al., 2024; Ni et al., 2020; Reiffel et al., 2015). Despite this growing mechanistic and therapeutic insight, no systematic framework currently exists to identify and evaluate optimal polytherapy strategies in a sex-specific manner, even as accumulating experimental and clinical evidence demonstrates sex-dependent differences exist in atrial electrophysiology, Ca^2+^ handling, and arrhythmia vulnerability (Giammarino et al., 2025; Herraiz-Martínez et al., 2021; Pecha et al., 2023; Ravens, 2018). To address this limitation, we built a regression-guided computational pipeline that leverages population-derived sensitivities to inform therapeutic design. While population-based modeling approaches are widely used to characterize variability and perform sensitivity analyses (Ni et al., 2018), here we use regression coefficients to systematically construct and prioritize sex-specific multi-target intervention strategies under explicit physiological constraints. This framework enables high-throughput evaluation of complex drug combinations while maintaining mechanistic insight. At the single-cell level, male and female forward simulations showed that multi-target strategies generally improved APD and Ca^2+^ handling and reduced arrhythmia vulnerability (**Fig. 4**). In 2D atrial tissue, sex-specific drug strategies preserved CV while increasing refractoriness, resulting in an overall increase in wavelength and a reduction in reentry vulnerability in both sexes (**Fig. 6**).

### Sex-Specific Mechanistic Insights

In males, the regression-guided pipeline revealed that most accepted interventions primarily consisted of Na^+^ and K^+^ channel blockers. This pattern is consistent with the need to prolong APD in cAF, where inhibiting repolarizing K^+^ currents supports recovery of APD and effective refractory period toward nSR levels. From our previous studies, (Herrera et al., 2023, 2025) inhibition of these currents is also associated with an increase in CaT_Amp_ and a decrease in the Alt_BCL-T_, indicating an overall antiarrhythmic effect. However, excessive K^+^ current inhibition can increase susceptibility to DADs, suggesting that concurrent Na^+^ channel inhibition reduces vulnerability to DAD-related triggered activity. Importantly, Na^+^ channel inhibition must be carefully balanced, as excessive block may reduce V_Max_ and CV and could promote conduction block and reentry, thereby potentially offsetting the antiarrhythmic effect of APD prolongation. Of note, a small number of regression-accepted interventions in cAF males increased DAD_BCL-T_, underscoring the need for forward simulations to evaluate emergent effects. Together, these results suggest that effective rhythm stabilization in males arises from coordinated modulation of Na^+^ and K^+^ currents to restore APD and CaT_Amp_ without enhancing triggered activity. Reentry was completely abolished when pharmacological modulation was combined with partial recovery of intercellular coupling, which further increased CV and wavelength, consistent with experimental evidence that shows improved gap-junctional coupling reduces AF inducibility . These effects are particularly relevant in structurally remodeled atria, where fibrotic disruption of cell-cell connectivity slows conduction and promotes reentry (Ma et al., 2021; Silva Cunha et al., 2024).

Therapeutic recovery in females also required incorporation of K^+^ channel inhibition to counteract cAF-induced shortening of APD and reduction of CaT_Amp_. However, a notable distinction from males was the higher prevalence of effective combinations that included inhibition of Ca^2+^ handling targets consistent with suppression of Ca^2+^ mediated DAD triggers through modulation of Ca^2+^ handling pathways (Faggioni et al., 2014). This observation aligns with our previous findings (Herrera et al., 2025; Ni et al., 2023; Zhang et al., 2024), that reducing Ca^2+^ influx or CaMKII-dependent LTCC phosphorylation decreases DAD susceptibility by limiting sarcoplasmic reticulum Ca^2+^ loading and the likelihood of spontaneous Ca^2+^ release during diastole. While such inhibition contributes to greater electrical stability, it can also depress CaT_Amp_, highlighting a trade-off between suppressing triggered activity and preserving contractile function, which likely contributes to the narrower range of effective inhibitor combinations observed in females compared with males.

### Polytherapy and Synergistic Effects

AF is characterized by the interaction of multiple arrhythmogenic mechanisms, including altered membrane excitability, impaired refractoriness, dysregulated Ca^2+^ handling, and conduction abnormalities (Nattel et al., 2008, 2020). Notably, early antiarrhythmic drugs for AF were developed without detailed knowledge of their molecular targets, yet achieved moderate clinical efficacy based on their effects on cardiac electrophysiology (Heijman et al., 2013, 2021; Katritsis & Camm, 1994; Zimetbaum, 2012). Many of these agents were later recognized to inhibit multiple ionic currents, revealing a degree of mechanistic pleiotropy not originally intended (Heijman et al., 2021). For example, amiodarone, is thought to exert broad antiarrhythmic effects through concurrent modulation of Na^+^, K^+^, and Ca^2+^ currents as well as adrenergic signaling, highlighting how coordinated multi-target modulation can influence atrial electrophysiology (Heijman et al., 2021; Mujović et al., 2020; Oryan et al., 2018). Furthermore, clinical and computational studies have increasingly supported this concept by demonstrating that coordinated modulation of multiple ionic targets, can produce synergistic improvements in atrial electrophysiological stability and more effectively suppress AF than single-channel block (Dasí et al., 2024; Ni et al., 2017, 2020; Reiffel et al., 2015). Consistent with these observations, our findings provide a mechanistic explanation for the clinical efficacy of multi-target antiarrhythmic therapy, demonstrating that coordinated inhibition can simultaneously stabilize multiple interacting AF mechanisms and that the specific targets required for this stabilization differ between males and females.

### Limitations and Future Work

Several limitations should be acknowledged. The tissue simulations assume spatially homogeneous ionic properties, whereas AF is characterized by pronounced regional heterogeneity in electrophysiology, Ca^2+^ handling, and gap-junction remodeling (Iwamiya et al., 2024), which may alter both DAD and triggered AP formation and reentrant dynamics. Similarly, recovery of cell-cell coupling was implemented as a uniform increase in diffusion, which serves as a proxy for improved gap-junctional communication or reduced fibrosis but does not explicitly represent heterogeneous connexin expression, anisotropic conduction, fiber orientation, or myocyte-fibroblast coupling. Incorporating spatially heterogeneous electrophysiological and structural properties, and remodeling patterns, will be important to determine the robustness of the identified sex-specific therapeutic strategies into more realistic substrates. Furthermore, extending our pipeline to anatomically realistic atrial geometries will allow assessing whether interventions that suppress triggered activity or eliminate reentry in simplified tissue remain effective in the presence of realistic boundaries, curvature, and conduction pathways. Additionally, the regression-guided pipeline focuses on steady-state pacing conditions and evaluates drug effects as fixed perturbations of ionic and Ca^2+^ handling parameters. Future studies will extend this framework to examine how dynamic, rate-dependent drug actions influence arrhythmia vulnerability across sexes (Ellinwood et al., 2017; Moreno et al., 2011; Morotti et al., 2016).

## Conclusions

This study establishes a sex-specific, regression-guided modeling pipeline for identifying and validating antiarrhythmic polytherapy strategies across cellular and tissue scales. By integrating regression-derived predictions with mechanistic single-cell and 2D tissue simulations, the pipeline reveals distinct therapeutic requirements in males and females with cAF, demonstrating that effective rhythm stabilization arises from sex-specific, multi-target modulation of ionic and Ca^2+^ handling pathways, and that interventions optimized at the cellular level remain effective in tissue when combined with preserved conduction. These findings support the use of sex-informed, multiscale modeling to guide the development of more effective and safer antiarrhythmic therapies against AF. Meaningful translation will require integrating the computational pipeline with experimental measurements of sex-specific drug responses in human atrial tissue, to validate the effects of specific pharmacological interventions on electrophysiological and Ca^2+^ handling effects and enhance the ability of multiscale models to guide mechanism-driven polytherapy design.

## Sources of Funding

American Heart Association Predoctoral Fellowship 24PRE1183427 (NTH)

American Heart Association Career Development Award 24CDA1269250 (CERS)

American Heart Association Career Development Award 24CDA1258695 (HN)

NIA Grant R03AG086695 (CERS, EG)

NHLBI Grant R01HL176651 (EG, DD, SM)

NHLBI Grants R01HL131517, R01HL170521, R01HL141214, and P01HL141084 (EG)

NHLBI Grants R00HL138160, R01HL171057, and R01HL171586 (SM)

NHLBI Grants R01HL136389, R01HL163277, R01HL160992, R01HL165704, and R01HL164838 (DD)

Deutsche Forschungsgemeinschaft Research Training Group 2989 (DD)

European Union large-scale network grant No. 965286 (MAESTRIA, DD)

